# X-ray 3D imaging of gene expression in whole-mount murine brain by microCT, implication for functional analysis of tRNA endonuclease 54 gene mutated in pontocerebellar hypoplasia

**DOI:** 10.1101/2019.12.19.882688

**Authors:** Olga Ermakova, Tiziana Orsini, Paolo Fruscoloni, Francesco Chiani, Alessia Gambadoro, Sabrina Putti, Maurizio Cirilli, Alessio Mezzi, Saulius Kaciulis, Marcello Raspa, Ferdinando Scavizzi, Glauco P. Tocchini-Valentini

## Abstract

Acquisition of detailed structural and molecular information from intact biological samples, while preserving cellular three-dimensional structures, still represents a challenge for biological studies aiming to unravel system functions. Here we describe a novel X-ray-based methodology for analysis of gene expression pattern in intact murine brain ex vivo by microCT. The method relays on detection of bromine molecules in the products of enzymatic reaction generated by the *β*-galactosidase (lacZ) gene reporter. To demonstrate the feasibility of the method, the analysis of the expression pattern of tRNA endonuclease 54 (Tsen54)-lacZ reporter gene in the whole-mount murine brain in semi-quantitative manner is performed. Mutations in Tsen54 gene causes pontocerebellar hypoplasia (PCH), severe neurodegenerative disorder with both mental and motor deficits. Comparing relative levels of Tsen54 gene expression, we have demonstrated that highest Tsen54 expression observed in anatomical brain substructures important for the normal motor and memory functions in mice. In the forebrain strong expression in perirhinal, retrosplenial and secondary motor areas was observed. In olfactory area Tsen54 is highly expressed in the nucleus of the lateral olfactory tract, anterior olfactory and bed nuclei, while in hypothalamus in lateral mammillary nucleus and preoptic area. In hindbrain Tsen54 is expressed in the reticular, cuneate and trigeminal nuclei of medulla, and in pontine gray of pons and in cerebellum, in the molecular and Purkinje cell layers. Delineating anatomical brain regions in which Tsen54 is strongly expressed will allow functionally address the role Tsen54 gene in normal physiology and in PCH disease.

**Significance Statement:** Characterization of gene expression pattern in the brain of model organisms is critical for unravelling the gene function in normal physiology and disease. It is performed by optical imaging of the two-dimensional brain sections which then assembled in volume images. Here we applied microCT platform, which allows three-dimensional imaging of non transparent samples, for analysis of gene expression. This method based on detection by X-ray the bromine molecules presented in the products generated by enzymatic activity of b-galactosidase reporter gene. With this method we identify anatomical brain substructures in which Tsen54 gene, mutated in pontocerebellar hypoplasia disease, is expressed.

## Introduction

Molecular phenotyping of gene expression by three-dimensional (3D) visualization of the biological samples is widely used to gain a comprehensive understanding of structure-function relationships in complex biological systems (1). Traditionally, this is achieved by histological dissection of the sample, visualization of gene expression patterns on consecutive sections by light microscopy and 3D digital assembly of the acquired two-dimensional (2D) information into a three-dimensional view (2, 3). Application of sophisticated reconstruction methods is required to obtain a 3D volume image from stacked 2D images without substantial shape and morphology loss (1, 4, 5, 6, 7).

Recently, innovative methodologies, such as two-photon microscopy, optical coherence tomography and confocal microscopy have been used for imaging of optically transparent tissues obtained by the clearing of whole mount samples, allowing for a direct 3D view of complex tissues (8, 9, 10,11, 12, 13, 14, 15, 16, 17,18). These techniques, while providing valuable information at cellular resolution, are technologically challenging and include lengthy sample preparation procedures. Moreover, the limited light penetration through the sample, which is still a flaw of these methods, often impose acquisitions of images through multiple planes and, consequently, the usage of dedicated software for the 3D reconstruction calculations. These methodological time consuming steps cause substantial delays that hinder their usage as routine analysis tools in laboratories.

Several international high-throughput projects aiming to the systematic analysis of gene expression were launched in recent years. One of the largest is the genome-wide gene expression profiling analysis in murine and human brains undertaken by the Allen Institute for Brain Research, which data are deposited and freely available to the scientific community (www.brain-map.org). As part of this effort, Mouse Brain Atlas project provides the neurological expression patterns of 20,000 genes assessed by *in situ* hybridization (ISH) of the histological mouse brain sections mapped on 2D murine anatomical reference brain atlas (19). Further, 3D representation of the ISH data is then achieved by digital assembly of 2D images across the brain performed by complex mathematical and statistical algorithms (20). Another high-throughput project is carried out by the International Mouse Phenotyping Consortium (IMPC), which has the goal to determine the gene functions by systematic physiological and molecular phenotyping of 20,000 knockout mouse strains (www.mousephenotype.org) (21). With the same aim, the International Mouse Knockout Consortium (IKMC) is generating knockout mouse strains for every protein-coding gene. The knockout first alleles produced by the IKMC are obtained by inserting a splice-acceptor containing *lacZ* reporter gene followed by a critical exon, common to all transcript variants, flanked by *LoxP* sites (https://www.beta.mousephenotype.org/about-ikmc-strategies) (22). Molecular phenotyping of these knockout animals includes a systematic analysis of gene expression patterns using *lacZ* reporter in several mouse tissues and in the brain specifically. The IKMC engineered animals as well as phenotyping data produced by IMPC consortium are freely available for international scientific community (23). Overall, the mentioned above resources provide an unprecedented wealth of information for understanding how the genes determine the correct structure-function relationships within complex biological systems and organisms.

Among the imaging techniques used for the analysis of biological samples in 3D, X-ray microtomography (microCT) has recently gained increased popularity and broad application in biomedical research. While the microCT imaging received extensive usage in biological studies of mineralized animal tissues (24, 25), analysis of soft tissues using X-ray is still a substantial challenge because of their low detectable adsorption in the X-ray spectrum. To circumvent this inherent constrain of the technique, several sample treatments with contrasting agents have been utilized to increase X-ray visualization of soft tissues. This expedient allowed to extend the inspection biological samples by microCT to several whole-mount *ex vivo* murine organs, brain blood vessels and murine embryos among others (26, 27, 28, 29, 30).

Tsen54 gene frequently mutated in patients affected by Pontocerebellar Hypoplasia (PCH). PCH is a heterogeneous group of autosomal recessive neurodegenerative disorders characterized by a wide diagnostic spectrum including the delay in cognitive and motor development, seizures and, death in early childhood (30). PCH subtypes morphologically characterized by severity of cerebellum and ventral pons hypoplasia, progressive microcephaly, neocortical atrophy (32).

Genetic analysis on PCH patients identified several pathological mutations in a small numbers of genes. The majority of PCH cases have been linked to mutations in genes important for tRNA metabolism. In particular, 60% for all genetically defined cases of PCH are caused by mutations in the genes encoding for protein subunits of the TSEN complex (31, 33, 34, 35, 36). The main function accomplished by the TSEN complex is to produce mature and functional tRNAs by removing an intron. The heteromeric TSEN complex is composed of four subunits: Tsen54, Tsen34, Tsen15 and Tsen2. The catalytic activity of this complex is devoted to Tsen2 and Tsen34 subunits, while Tsen54 and Tsen15 are just structural but critical for correct orientation of the catalytic sites within the complex subunits (37). Recessive mutations in all four subunits of the TSEN complex have been identified in PCH patients with different degree of severity and are classified as PCH types 2A-C, 4 and 5 (33, 34, 38, 35, 39). It has been observed that all homozygous Tsen54 mutation carriers had a phenotype consistent with PCH2A subtype (33). Most of the patients carrying the Tsen54 pathogenic mutations exhibit abnormal muscle tone since earlier months after birth, central vision impairment, epileptic seizures, and severe psychomotor retardation (31).

It human developing brain Tsen54 gene is highly expressed in the telencephalon and metencephalon at 8 weeks of gestation (40). The telencephalon gives rise to the cerebral cortex and basal ganglia within the cerebral hemispheres. The metencephalon region of the developing brain eventually forms the pons and cerebellum. Indeed, at 23 weeks of gestation a strong and specific expression of TSEN54 in developing human cerebellar neurons has been reported (38). While the Tsen54 mRNA has been extracted and detected in human cortex and cerebellum (https://www.genecards.org/cgi-bin/carddisp.pl?gene=Tsen54 and www.human.brain-map.org), the histological characterization of the Tsen54 expression pattern in the adult brain was not performed. In model organisms analysis of Tsen54 gene expression demonstrated that Tsen54 gene expressed ubiquitously in developing zebrafish (40). In murine brain the *in situ* hybridization data contained in the Allen Brain Atlas database, suggest very low, difficult to interpret signal (https://mouse.brain-map.org/experiment/69352788).

In this manuscript, we describe a three-dimensional expression analysis of the *Tsen54-lacZ* reporter gene by X-ray imaging in whole-mount *ex vivo* murine brains. We accomplished such mapping by applying our in house developed method, which we previously successfully used for the analysis of *lacZ* reporter gene expression in ependymal cells of the murine brain embedded in paraffin (41). Our custom protocol relays on the *in situ* detection of the *lacZ* reporter gene enzymatic activity, which give raise to dense products detectable by X-ray imaging directly in whole-mount brains. Because the basal endogenous contrast of the rest (i.e. non-expressing *lacZ* regions) of paraffin embedded murine brain is unaffected, our method allows for the visualization of anatomical brain substructures where the *lacZ* reporter gene is active. Because the source of such increased in X-ray absorption was still matter of debate, we also analysed what component of the reaction is responsible for this observation. Our data suggest that the bromine atoms within the *lacZ*/*X-gal* conversion precipitate *in situ* are the main X-ray contrasting agent.

We also performed a density-based semi-quantitative estimation of *Tsen54-lacZ* gene expression in adult mouse brain, identifying anatomical substructures with high levels of *Tsen54-lacZ* expression. 3D virtual models of *Tsen54-lacZ* expression were build based on this analysis. These findings will allow to focus further studies to the identified brain with the aim to understand the function of *Tsen54* gene in these brain structures in the normal development and mechanisms leading to PCH disorder.

## Results

### Three-dimensional visualization of the Tsen54-lacZ reporter gene expression in the whole-mount mouse brain

In order to demonstrate the feasibility of microCT imaging for successful 3D studies of gene expression, we performed several scanning arrays of whole-mount mouse brains of the *Tsen54-lacZ* (IKMC-*Tsen54^Tm1b/+^* line) reporter animals. To analyse the expression of Tsen54 gene we took advantage of the presence of *lacZ* reporter gene. Whole mouse brains were assayed *for X-gal*/*FeCN β-galactosidase* activity, using *X-gal* (*5-bromo-4-chloro-3-indolyl β-D galactopyranoside*) as a substrate for enzymatic detection of *β-galactosidase* expression in combination with potassium ferri- and ferro-cyanide (*X-gal*/*FeCN* protocol) as suggested by IMPC consortium (www.kompphenotype.org) (42). In our experiments, the blue precipitate product of *X-gal*/*FeCN* staining is strong and visible in whole brain preparations from the*Tsen54-lacZ* mice, but not in the wild type controls (Figure 1B, C).

**Figure 1.**
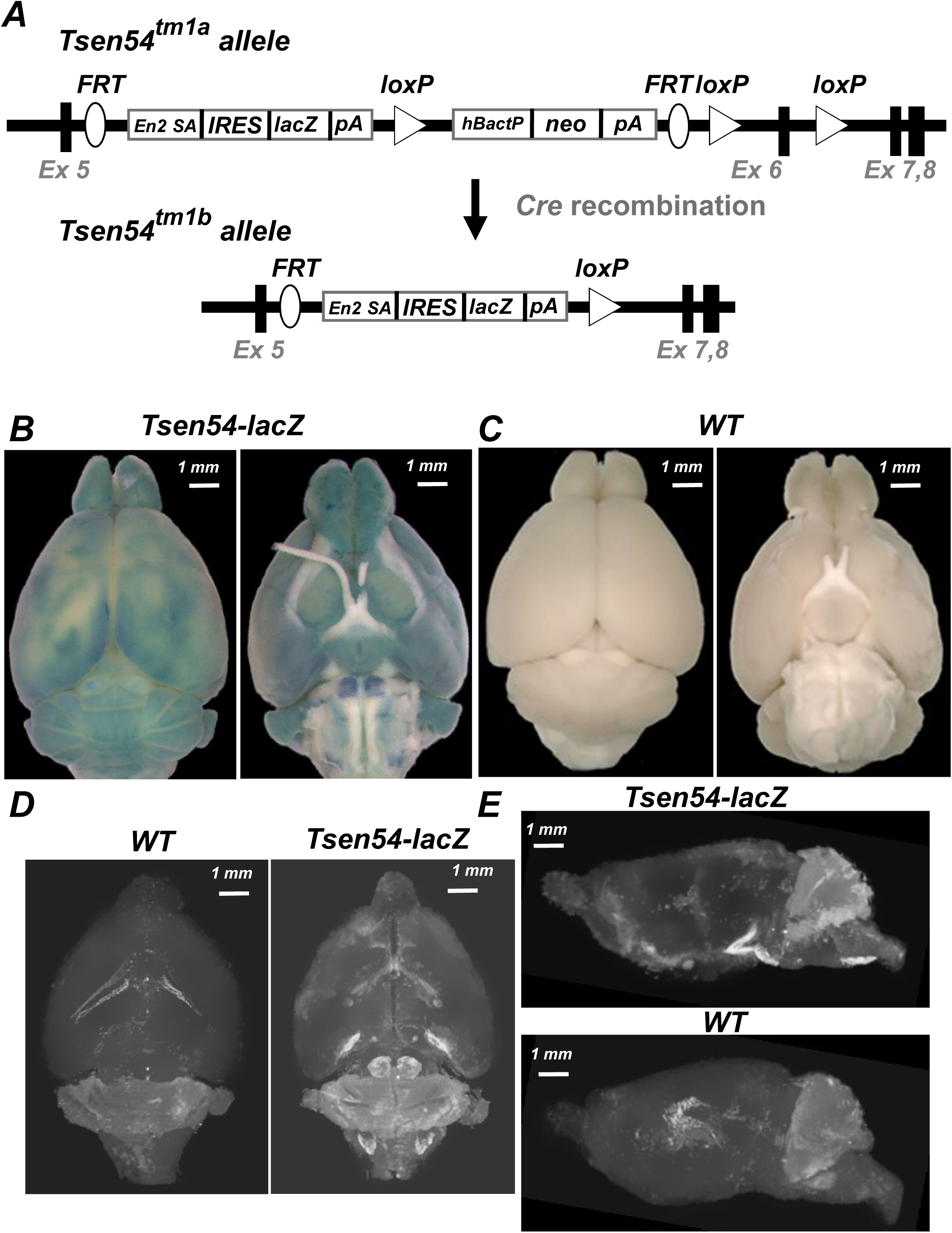
Three-dimensional imaging of Tsen54-lacZ expression in intact adult mouse brain by microCT. **A.** Schematic representation of the *Tsen54^Tm1a^* allele. *lacZ* reporter gene is inserted into intronic locus, following exon5, replacing exons 6 of the *Tsen54* gene. *Cre*-mediated recombination removes exon6 and neo targeting cassette from genomic *Tsen54^Tm1a^* gene locus and convert *Tsen54*^Tm1b^ allele into *Tsen54^Tm1b^* allele. The *Tsen54^Tm1b^* allele is a knockout of *Tsen54* gene and is a *lacZ* reporter allele of *Tsen54* gene expression. *Ex*-exon; *En2A SA*-splice acceptor site; *IRES*-internal ribosomal entry site; *lacZ* - bacterial *β-galactosidase* reporter gene; *pA*-poly A; *hBactP*-human β-actin promoter; *neo*-neomycin resistance gene; *FRT-FLP* recombination sites and *LoxP* – *Cre* recombination sites are indicated. ***B.*** Dorsal and ventral views of whole mouse brain from *Tsen54-lacZ* reporter/knock-in allele (IKMC*-Tsen54*^Tm1b^ allele) stained with *X-gal/FeCN* visualized by stereomicroscope; ***C.*** Dorsal and ventral views of littermate control whole mouse brain stained with *X-gal/FeCN* visualized by stereomicroscope***; D.*** Representative Maximum Intensity Projection (MIP) microCT images scanned with the resolution 5 μm/voxel of whole brain from *Tsen54-lacZ* (N=3) and *WT* (N=3) animals. The regions of increased density (white) correspond to precipitated product of *X-gal* substrate converted b*y β-galactosidase. **E.*** Sagittal view of MIP whole brains images represented in panel ***D***.

To examine, whether the deposited products of *β-galactosidase* reaction provide sufficient X-ray density to mark the regions of *lacZ* reporter expression, we performed microCT scans of the whole-mount *X-gal*/*FeCN* stained, dehydrated and paraffin embedded brains from *Tsen54-lacZ* animals. MicroCT imaging detected the regions of increased density in the *Tsen54-lacZ* brains, which were not observed in wild type brains (Figure 1D). Strikingly, our Maximum Intensity Projection (MIP) of microCT images showed remarkable overlaps with images we obtained by light stereomicroscopy of the mouse brains (Figure 1B). MIP of volume images also allowed for throughout inspection of the sample, both in frontal and sagittal 2D views (Figure 1D, E). To fully appreciate the 3D views of these results we provide the movies of the *Tsen54-lacZ* and wild type brains as downloadable files (***Movies 1, 2***).

To further analyse the nature of the detected by X-ray contrast differences between *Tsen54-lacZ* and wild type brains volumes, we compared respective 2D microCT derived sections from wild type and *Tsen54-lacZ* brains. Since, a volume microCT image is a three-dimensional matrix of brightness values expressed in voxel, digitally reconstructed from multiple 20 μm 2D sections, it can be digitally dissected and presented as a digital 2D sections (Figure 2). Comparing 2D sections from the same brain regions of the *Tsen54-lacZ* and wild type, the areas of the high contrasts were observed in *Tsen54-lacZ* sections which were absent from the wild type samples (Figure 2A).

**Figure 2.**
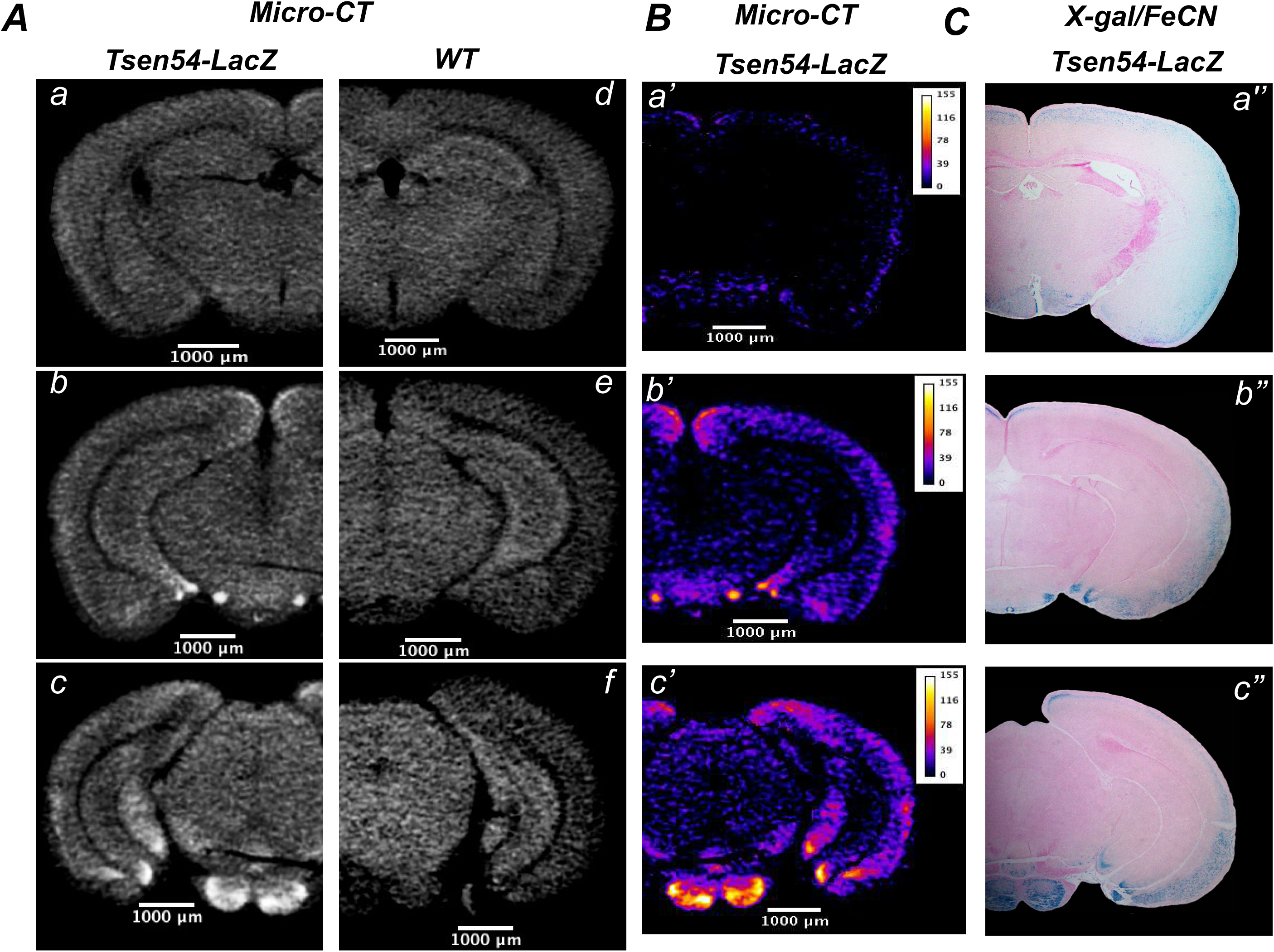
2D microCT-derived sections of forebrain reveal high-density X-ray patterns in good correspondence with chromogenic product of lacZ reporter detected by histological analysis. *A.* Two-dimensional MicroCT-derived sections from forebrain of *Tsen54-lacZ* and wild type animals whole-mount stained with *X-gal/FeCN:* from *Tsen54-lacZ* brain (a-c); corresponding sections from wild type brain(d-f). *B.* Segmentation analysis of the 2D microCT-derived sections of *Tsen54-lacZ*;(a’-c’). *C.* Best corresponding histological sections from the *Tsen54-lacZ* brain whole-mount stained with *X-gal*/*FeCN* and sectioned for histological analysis (a’’-c’’).

To define the regions, in which X-ray reveals increase in densities above the background computationally we performed a segmentation analysis of 2D sections derived from *Tsen54-lacZ* brains. The numerical value computed for each voxel of microCT image is a linear X-ray attenuation coefficient expressed as a gray value, at the corresponding point in the sample volume (43). We inferred that reconstructed X-ray absorption values at each voxel location in the acquired images of experimental sample is the sum: 1) of the brightness derived from deposited products of *lacZ* enzymatic reaction and 2) endogenous values of the dehydrated tissue. Therefore, the expression level of the *lacZ* reporter could be determined after subtraction for the background values calculated using microCT images derived from wild type samples. We performed this calculation using ImageJ program. Since endogenous grey values differ between the different anatomical substructures even in murine wild type brain we delineated following regions in murine forebrain: cortex, mid-brain, hippocampus and amygdala. This operation was feasible because the anatomical structures can be easily defined on microCT derived 2D sections (Figure 2A). Calculated from wild type sections, mean and one standard deviation of gray scale values for each region were then subtracted from the corresponding brain region of experimental *Tsen54-lacZ* brain section. The LUT function from 0-155 gray values was set for the resulting images using ImageJ program (Figure 2B).

To illustrate that the X-ray opaqueness is indeed produced by *β-galactosidase* reaction products *in situ*, we compared segmented microCT images and corresponding histologically analysed sections. To this purpose, *X-gal*/*FeCN* stained and paraffin embedded brains after microCT imaging were sectioned and reanalysed by light microscopy (Figure 2C). Overall, a good agreement between regions with elevated densities detected by microCT and *the X-gal*/*FeCN* chromogenic precipitates, product of *lacZ* reaction, was observed when the forebrain 2D sections were analysed (Figure 2B,C). Importantly, histological analysis of control samples did not reveal any chromogenic product of *lacZ* activity in the brains of the wild type littermates (data not shown) and this result is in agreement with previously published data (42).

We then compared 2D microCT-derived images of the hindbrain region of *Tsen54-lacZ* and wild type brains (Figure 3). The hindbrain was subdivided on the two regions: 1) cerebellar lobes and 2) medulla and pons. This sectioning was necessary because of the substantial divergence in the medium gray values of these hindbrain substructures even in wild type brains. Sum of the mean and standard deviation calculated for each region separately were then subtracted from the *Tsen54-lacZ* respective subregions of hindbrain (Figure 3B). The intensity of the remaining signal then was calibrated, using LUT function of the ImageJ software. Further, we compared the segmented microCT sections to the best anatomically corresponding histological section of the hindbrain. This comparison demonstrated that regions with the high X-ray densities are in good agreement with the regions in which chromogenic *lacZ* product was detected also by histological analysis.

**Figure 3.**
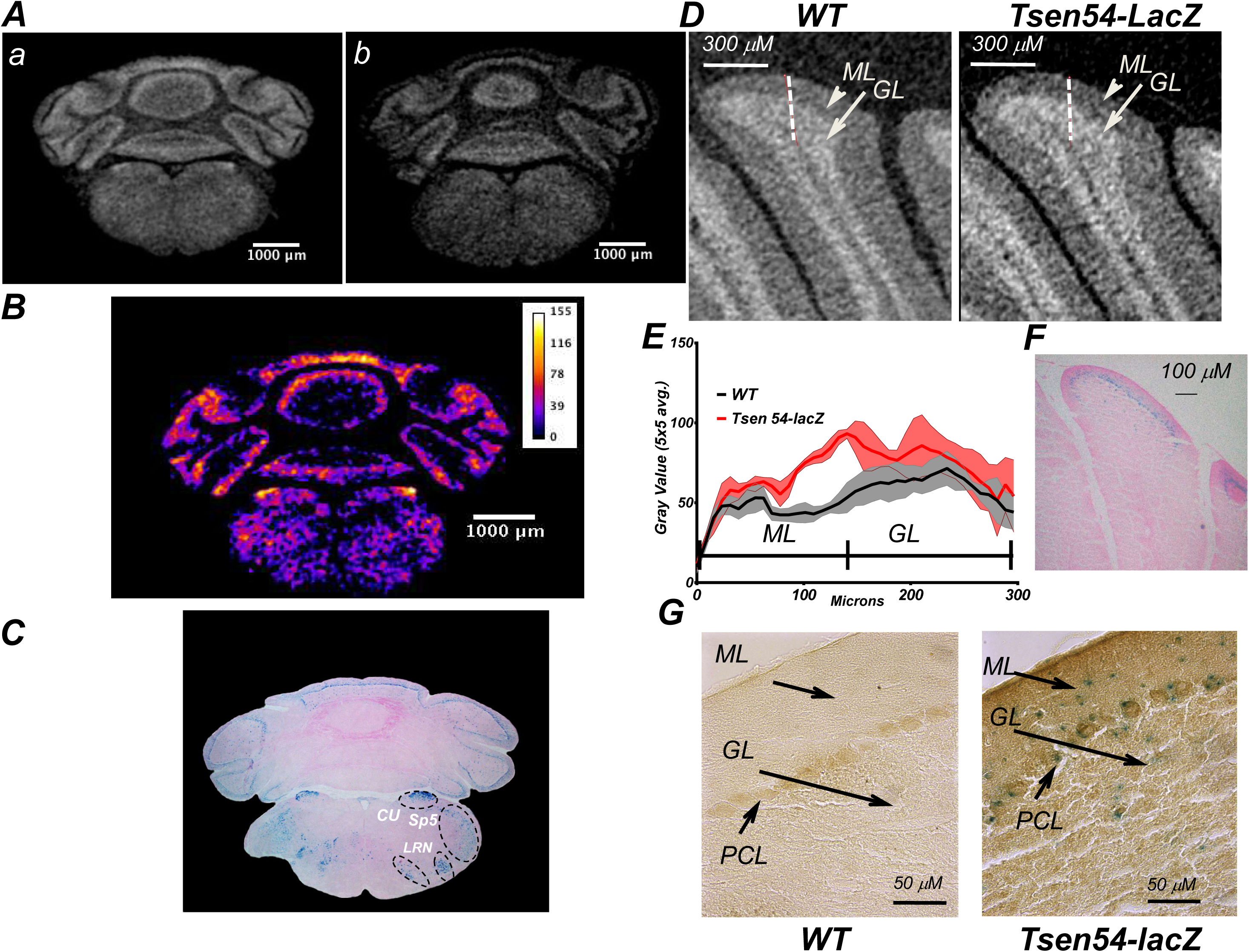
2D MicroCT and histological analysis of Tsen54-lacZ reporter expression in murine hindbrain. ***A.*** Representative 2D dimensional microCT-derived sections from hindbrain region: *Tsen54-lacZ* (***a***) and wild type (***b***) brains. ***B.*** Segmentation analysis of the 2D microCT hindbrain section, using ImageJ program. ***C***. Histological section of hindbrain from the whole-mount *X-gal/FeCN* stained *Tsen54-lacZ* brain. The regions with the highest expression in medulla are indicated: *CU*-cuneate nucleus; *Sp5*-spinal nucelus of trigeminal; *LRN*-lateral reticular nucleus. ***D.*** 2D microCT acquired image of the simple cerebellar lobule from the *Tsen54-lacZ* and wild type animals at 4.9 μm/voxel resolution. The punctate lines indicate the regions selected for density quantification. Arrows indicate the structures of cerebellum: *ML*-Molecular Layer and *GL*-Granular Layer. ***E.*** Average of brightness values, corrected for background, from *Tsen54-lacZ* (N=3; males) and wild type (N=3;males) cerebellar lobule, were plotted as a function of distance. ***F.*** Histological analysis of the simple cerebellar lobule of cerebellum from the whole-mount *X-gal/FeCN* stained brains. ***G.*** Immunohistochemical analysis of the *lacZ* expression with anti *b-galactosidase* antibodies of the *Tsen54-lacZ* and wild type cerebellar lobes: *ML*-Molecular Layer, *PCL*-Purkinje Cell Layer and *GL*-Granular Layer are indicated by the arrows.

Analysing hindbrain region we observed that the cerebellar lobes characterized by high contrast values. In addition, two main layers composing cerebellar lobes are distinguishable by a very different contrast, such that molecular and granular layers of cerebellum can be easily visualized and distinguished on the 2D microCT derived sections (Figure 3D). The highest contrast values were detected in the cerebellar granular layer, both in wild type and *Tsen54-lacZ* brains. The high endogenous background contrast complicates calculation of the signal derived specifically from the product of the *X-gal*/*β-galactosidase* reaction.

To demonstrate that we can digitally distinguish the regions of *lacZ* reporter activity from the non-expressing ones, we focused on analysis of molecular and granular layers of simple lobe in *Tsen54-lacZ* and wild type cerebellum (Figure 3D). This analysis was performed using Bruker CTAN program. We quantified the brightness (gray values) within the region of cerebellar simple lobe in *Tsen54-lacZ* reporter and wild type animals after background correction. The background value was calculated averaging pixels values over 300 μm length derived from paraffin (correction for the paraffin embedding). The average background value was then subtracted from microCT measured gray value for every pixel. The pixel values were measured over 300 mm length, three parallel measurements were performed with the 100 mm interval and mean was calculated for three adjacent pixels and plotted (Figure 3E). The graphical representation demonstrated an increased density in *Tsen54-lacZ* reporter animals. The maximum range of calculated densities between wild type and *Tsen54-lacZ* is observed at the boundary between molecular and granular layers (Figure 3E). This observation is in agreement with the histological analysis. Optical analysis showed that the chromogenic product of *β-galactosidase* reaction was observed within internal molecular layer and a very strong signal was observed between molecular and granular layers (Figure 3F). We noticed, however, that while visual correspondence between the chromogenic products of b*-galactosidase* reaction with observed increase in density at the boundary between molecular and granular layers are in good agreement, the distances between microCT obtained and histological measurements did not perfectly overlap. Such differences could be due to mechanical distortion of the cerebellar layers during sectioning for histological analysis, which could cause the decrease of cerebellar layer volumes.

Another observation that we made is that the microCT profile indicated increase in X-ray density both in molecular and in granular layers of *Tsen54-lacZ* cerebellum, with the latter showing absence of *lacZ* staining. We hypothesized that *β-galactosidase* was expressed in granular layer as well, albeit at levels that are not high enough to produce a suitable amount of optically detectable chromogenic product after X-gal*/FeCN* staining. We therefore tried to increase the sensitivity of *β-galactosidase* detection by using *anti-β-galactosidase* antibodies and immunohistochemical analysis of the sections, previously stained with *X-gal/FeCN*. As expected, we observed that, in the *Tsen54-lacZ* samples, the *X-gal* stained cells are also positive for *β-galactosidase* protein. However the subset of the granular layer cells was negative for the visible product of enzymatic reaction, but positive for *β-galactosidase* immunoassay (Figure 3G).

The immunohistochemistry demonstrated that in the Tsen54-lacZ animals the *β-galactosidase* is expressed in all cerebellar structures, such as molecular, Purkinje cell and granular cell layers albeit at different levels (Figure 3G). Granular cells showed very low expression of *Tsen54-lacZ*, detected only by immunohistochemistry assay, while the stellate and basket cells of molecular as well as Purkinje cells showed a higher level of expression that could be detected also *by X-gal* staining. Importantly, we observed that the Purkinje cells of wild type animals are positive for endogenous *β-galactosidase* protein. These data suggest that the background detected in cerebellum of wild type animals could be partially explained by the endogenous β−*galactosidase* activity in Purkinje cells of the cerebellum (Figure 3G).

Thus, microCT data overall are in good agreement with the histological analysis. We therefore concluded that the X-ray detected densities in the *lacZ* positive samples specifically increased in regions where *β-galactosidase* activity is also present. In addition, we observed, that in some cells the activity of *β-galactosidase* was not abundant enough to convert the *X-gal* substrate into chromogenic products, but it can be detected by X-ray imaging.

In conclusion, our results demonstrate that the biochemical *lacZ*/X-gal/*FeCN* reaction produces a contrasting agent detectable by X-ray imaging, allowing for visualization of the *lacZ* gene reporter activity directly in whole-mount brains using microCT settings.

### Semiquantitative analysis of the Tsen54-lacZ gene expression in murine brain

In order to explore the possibility of a quantitative volumetric evaluation of the *lacZ* reporter expression, we compared the microCT-derived density histograms obtained from the experimental and control brain samples. The values in Hounsfield Units (HU) were plotted against voxel frequency of the *Tsen54-lacZ* and wild type brains datasets (Figure 4A). The histograms showed two peaks: one peak from 0-500HU and a second from 500 to max HU values. Using CTAN automatic threshold calculation with OTSU method (Bruker) we established that the peak at 0-500 HU range is caused by the X-ray absorption of the paraffin in which the brain samples were embedded, while the second peak corresponds to the brain tissues. A statistically significant shift in the HU densities versus maximum frequency values is observed when *Tsen54-lacZ* histogram was compared to the histogram of wild type brains.

**Figure 4.**
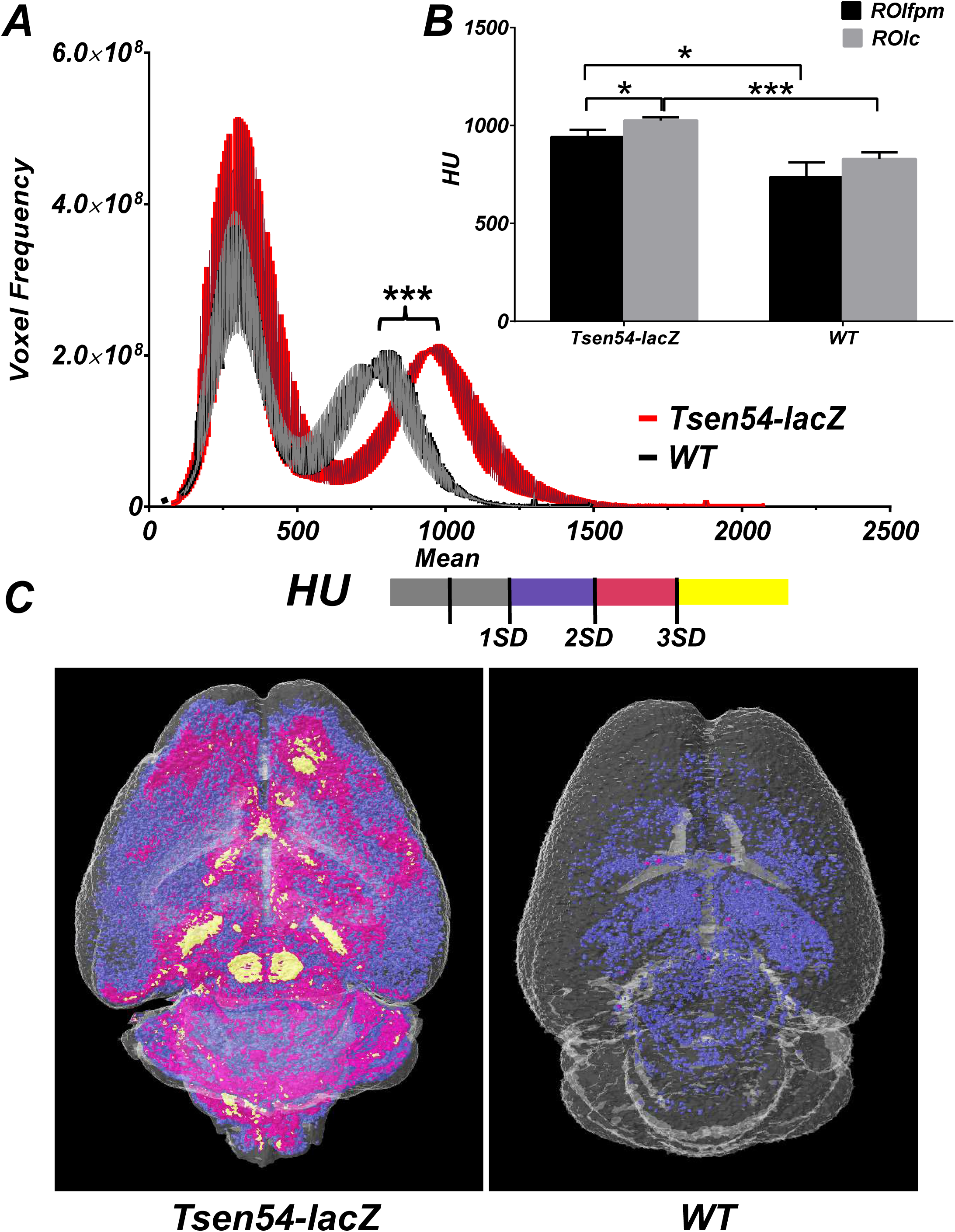
Relative quantification of Tsen54-lacZ gene expression by microCT analysis. ***A.*** Average histograms from *Tsen54-lacZ* and wild type brains stained by *X-gal/FeCN* method and imaged by microCT. Voxel frequencies of raw reconstructed sections of the brains were plotted against Hounsfield unit values (mean± SEM; *Tsen54-lacZ*, N=3;males; and WT, N=3;males; Mann-Whitney one-tailed statistical test; *** P < 0.001). ***B.*** Comparison of the Hounsfield densities using defined region of interests (ROI) between *Tsen54-lacZ* and wild type brains (mean± SEM; Tsen54-lacZ (N=3); WT (N=3); Two-tailed unpaired t-test; * P < 0.05; *** P < 0.001 *ROIfpm*-(ROIforebrain_pons_medulla); *ROIc*- (ROI_cerebellum). ***C.*** Comparative analysis of microCT derived densities in *Tsen54-lacZ* and wild type brains. Mean and standard deviation were calculated using CTAN (Bruker) program, using *Tsen54-lacZ* brains. Hounsfiled unit values inside of the mean +1standart deviation (1SD) were defined as a background, while values outside of this range were colour coded: 1SD to 2SD-blue; 2SD-3SD-magenta; 3SD and up to maximum value-yellow; The same Hounsfield value ranges were applied to analysis of the wild type brains.

To perform semi-quantitative analysis of reporter gene expression level, we calculated the means and standard deviations for the experimental and control samples using CTAN Bruker software. In order to accomplish this task, we selected two regions of interests (ROI): one ROIfmp delineating forebrain with medulla and pons and a second ROIc defining cerebellar vermis and hemispheres. Cerebellum were separated from the forebrain, medulla and pons regions because endogenous elevated density of the granular cells caused a skewed measurements towards the higher average densities and were interfering with the correct analysis of the brain regions outside of cerebellum. We therefore performed calculation of the two ROI regions separately. Mean HU densities of ROIfmp and cerebellum ROIc were then compared (Figure 4B). As a result, we observed statistically significant differences in average densities between ROIfmp and ROIc in *Tsen54-lacZ* brains. In wild type brains, the same trend was observed albeit the statistically significant differences were not reached. Importantly, statistically significant differences were also observed when ROIfmp and ROIc of *Tsen54-lacZ* when compared to the relevant ROI’s of the wild type brains. This observation is explained by higher X-ray attenuation due to the *lacZ*/X-gal/*FeCN* reaction products deposited *in situ* in the *Tsen54-lacZ* brains.

We then build a segmented 3D model of *Tsen54-lacZ* expression, using CTAN Bruker software. The HU values below the sum of mean and one standard deviation (SD) calculated for *Tsen54-lacZ* brain were empirically set as a background threshold. The HU values above the threshold were subdivided in three grades: from 1SD-2SD (lower corresponding density values); 2SD-3SD (intermediate) and from 3SD to max (higher density values). This HU values were implemented for segmentation analysis of both *Tsen54-lacZ* and wild type brains (Figure 2C). Using this approach, we were able to perform a relative quantification of the *Tsen54-lacZ* reporter gene expression levels (***Movie 3***).

The above results allowed us to produce a volumetric digital mapping of the *Tsen54* gene expression in the mouse brain. We used brain Brain Explorer2 software (Allen Brain Common Coordinate Network) to get a virtual 3D mouse brain showing only the regions in which the expression of the *Tsen54-lacZ* reporter was detected at elevated levels (Table 1, ***Movie 4***). The comparison between the images from microCT volume and those reconstructed using Brain Explorer2 software (Allen Brain Common Coordinate Network) allows to appreciate how the presented microCT imaging analysis can be instrumental for identification of the brain structures in which the expression of a gene, *Tsen54* in our case, occurs.

**Table 1:**
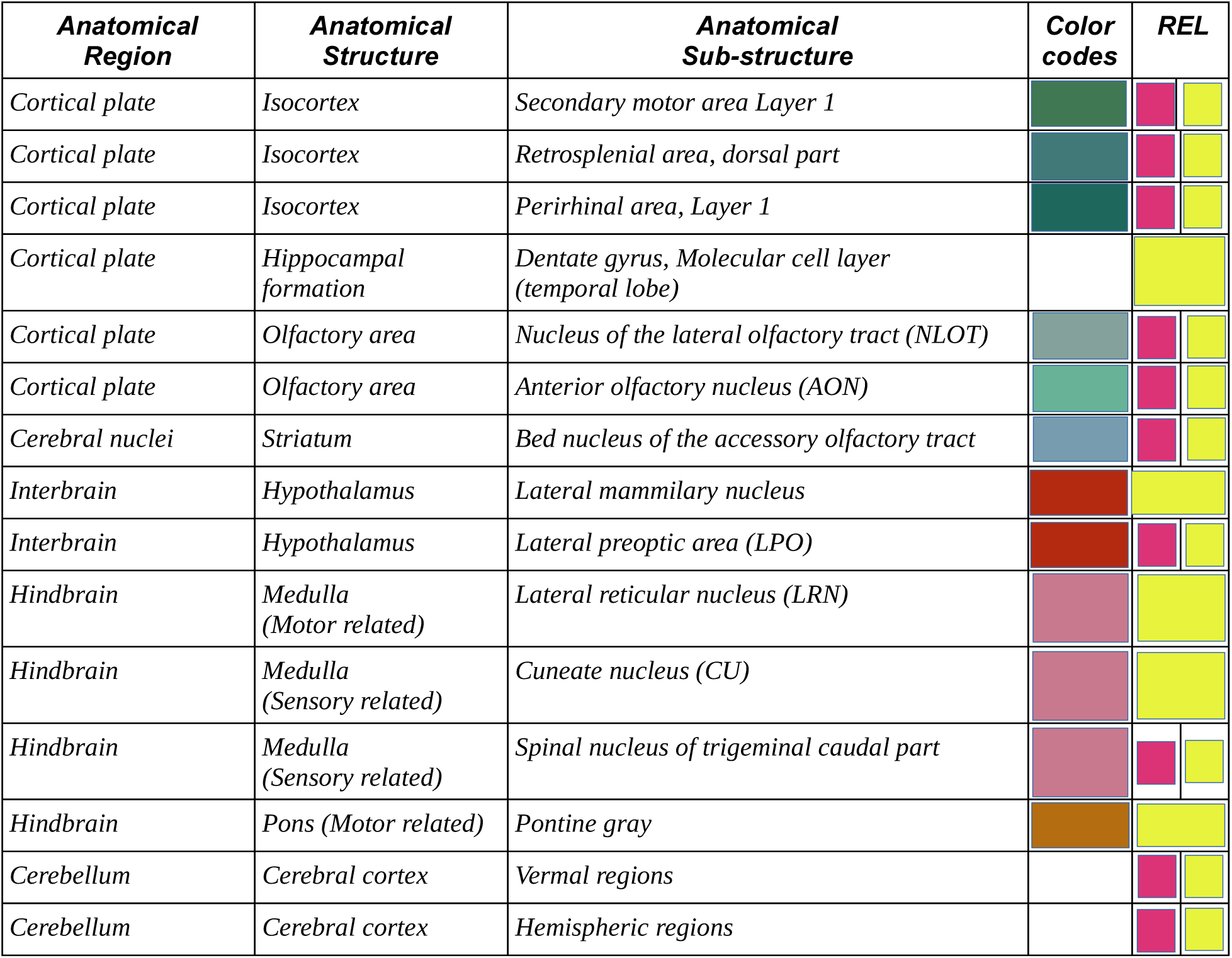
Summary of the anatomical substructures characterized by high expression of Tsen54-lacZ reporter gene expression. Anatomical brain structures and substructures in which Tsen54 gene is highly expressed are listed. The colour code for the indicated regions is extrapolated from A. Brain Atlas BrainExplorer 2 software. REL-relative expression level of the reporter gene. Yellow indicate highest expression level; magenta-intermediate level of the gene expression.

### Bromine in the β-galactosidase/X-gal reaction product is an X-ray contrasting atom

The methodology presented here couples the chemical detection of *lacZ* enzymatic activity, that converts substrate *X-gal* in chromogenic 5,5’-dibromo-4,4’-dichloro-indigo in presence of ferri- and ferro-cyanide, with *in situ* molecular signal within the mouse brain differentially increasing X-ray attenuation coefficient. Therefore, we aimed to determine molecular substrate detected by X-ray imaging. The biochemistry of the *β-galactosidase*/X-gal reaction has been extensively studied since the early 1950’s (44, 45). In brief, under proper conditions, *β-galactosidase* breaks the β-D-glycosidic linkage of *X-gal* releasing a soluble, colourless indolyl monomer. Subsequently, two liberated indolyl moieties can form a dimer, if a proper acceptor to which they can transfer an electron is present in solution (Figure 5A). The ferric and ferrous ions are included in most *X-gal* reaction buffers exactly to provide this function (46). The above chemical events produce a very stable and insoluble organic blue colour compound, which is an optically detectable *lacZ* reporter at the site of the reaction. We therefore asked, which compound, among the *lacZ*/*X-gal* reaction products, is able adsorb the X-rays and acts as an X-ray reporter in our experiments. The first hypothesis was that the presence of the electron transfer within the ferric/ferrous mix (*FeCN*) added to the *X-gal* reaction can lead to insoluble iron containing products, like FeIII, [FeIIIFeII(CN)6]3, similarly to the reaction which occurs during Prussian blue synthesis (47). We asked if possible formation of FeIII during the reaction could create a substrate that increase the X-ray attenuation coefficient at the sites of *β-galactosidase* enzymatic reaction.

**Figure 5.**
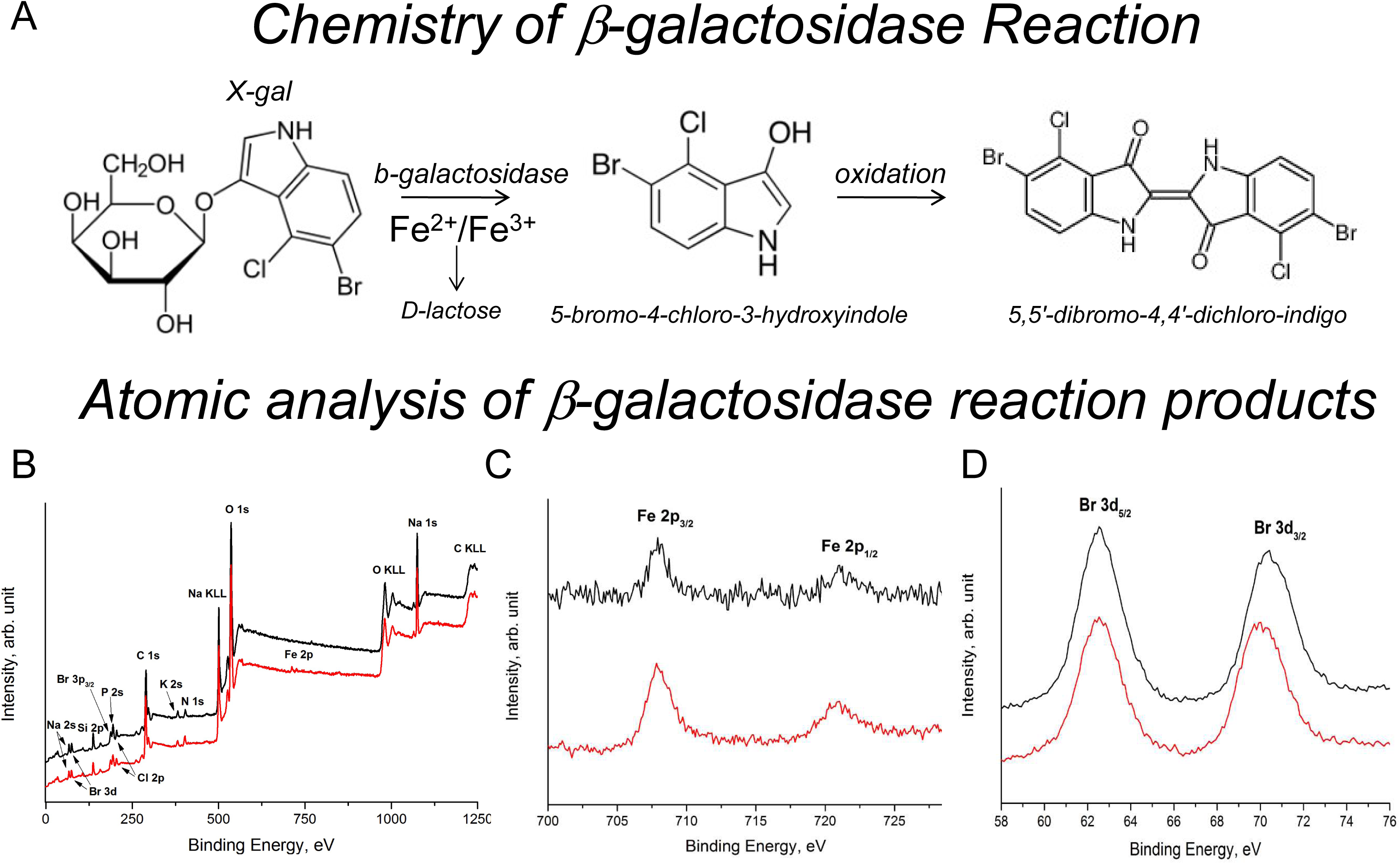
XPS analysis of the b-galactosidase enzymatic reaction products. ***A.*** Schematic representation of the b-galactosiadase reaction *in vitro*. ***B-D.*** X-ray photoemission spectroscopy (XPS) analysis of the *b-galactosidase* reaction products: black lines-reaction *b-galactosidase* with *X-gal/FeCN* substrate; red lines – *X-gal/FeCN* substrate only; ***B***. Complete survey spectra of the *b-galctosidase* reaction; ***C,D*** regions of main photoemission spectra Fe 2p (***C***) and Br 3d (***D***) peaks; black lines-reaction *b-galactosidase* with *X-gal/FeCN* substrate.red lines -substrate only;

To determine the molecules that could generate the high-contrast signal, we examined the atomic composition of the *β-galactosidase*/*X-gal/FeCN* reaction products using X-ray photoemission spectroscopy (XPS). The atomic components of the reaction, which were analysed *in vitro* with and without pure *β-galactosidase* in the presence of *X-gal*/*FeCN* substrate (Figure 5B-D). The XPS measurements excluded the presence of FeIII in the *β-galactosidase* reaction, because the Fe 2p_3/2_ peak at BE = 708.7 eV (Figure 5C) and the absence of satellite peaks at higher BE (lying at ∼ 6 eV or ∼ 8eV from the main component), are characteristic for FeII (48). Indeed no differences in atomic composition between reaction in the presence of *β-galactosidase* and control (substrate only) were detected.

To further confirm this finding, we performed a *Tsen54-lacZ* staining using tetranitroblue tetrazolium chloride as an electron acceptor instead of ferri- and ferro-cyanide (*FeCN*) solution. Whole-mount brains were stained with *X-gal*/tetrazolium, dehydrated, embedded in paraffin and imaged by microCT. (Figure 6). The *Tsen54-lacZ/X-gal*/tetrazolium stained brains were segmented, using the protocol described previously where Hounsfiled values below mean and 1SD were considered as a background and HU values above were defined in three steps: lowest range from 1SD-2SD; intermediate range from 2SD-3SD and highest above 3SD of HU. The comparison between the X-gal/tetrazolium and X-gal/FeCN methods demonstrated similar patterns of differential X-ray detected densities distributions. This finding holds true both for the regions in which *Tsen54-lacZ* gene expression were detected, and for the relative intensities of the reporter gene expression levels in specific brain areas (***Movie 3*** and ***Movie 5***). Importantly, the reproducible expression pattern of *Tsen54-lacZ* reporter imaged by microCT was obtained irrespective of which staining method and segmentation protocols were applied (Figure 2 and Figure 6***, Movies 3 and 5***). We therefore, ascertained that the bromine atoms contained in the chromogenic product of *X-gal*/*lacZ* reaction are responsible for the strong X-ray contrast observed *in situ* in brain tissues expressing *lacZ* reporter. Indeed, bromine has a relatively high atomic number (Z = 35) which generates a high-contrast enhancement for X-rays (49). The final chromogenic product of the *β-galactosidase* reaction contains two bromine molecules creating a marked increase in X-ray contrast enhancement (Figure 5A). To conclude, we demonstrated here that *in situ lacZ*/*X-gal* reaction products in whole-mount murine brain provide a high-contrast enhancement measurable by X-ray microCT imaging. Using this method, we were able to characterize the 3D gene expression pattern of *Tsen54-lacZ* reporter expression and to compare its relative levels in specific regions of murine brain, describing anatomical structures characterized by elevated level of *Tsen54* expression.

**Figure 6.**
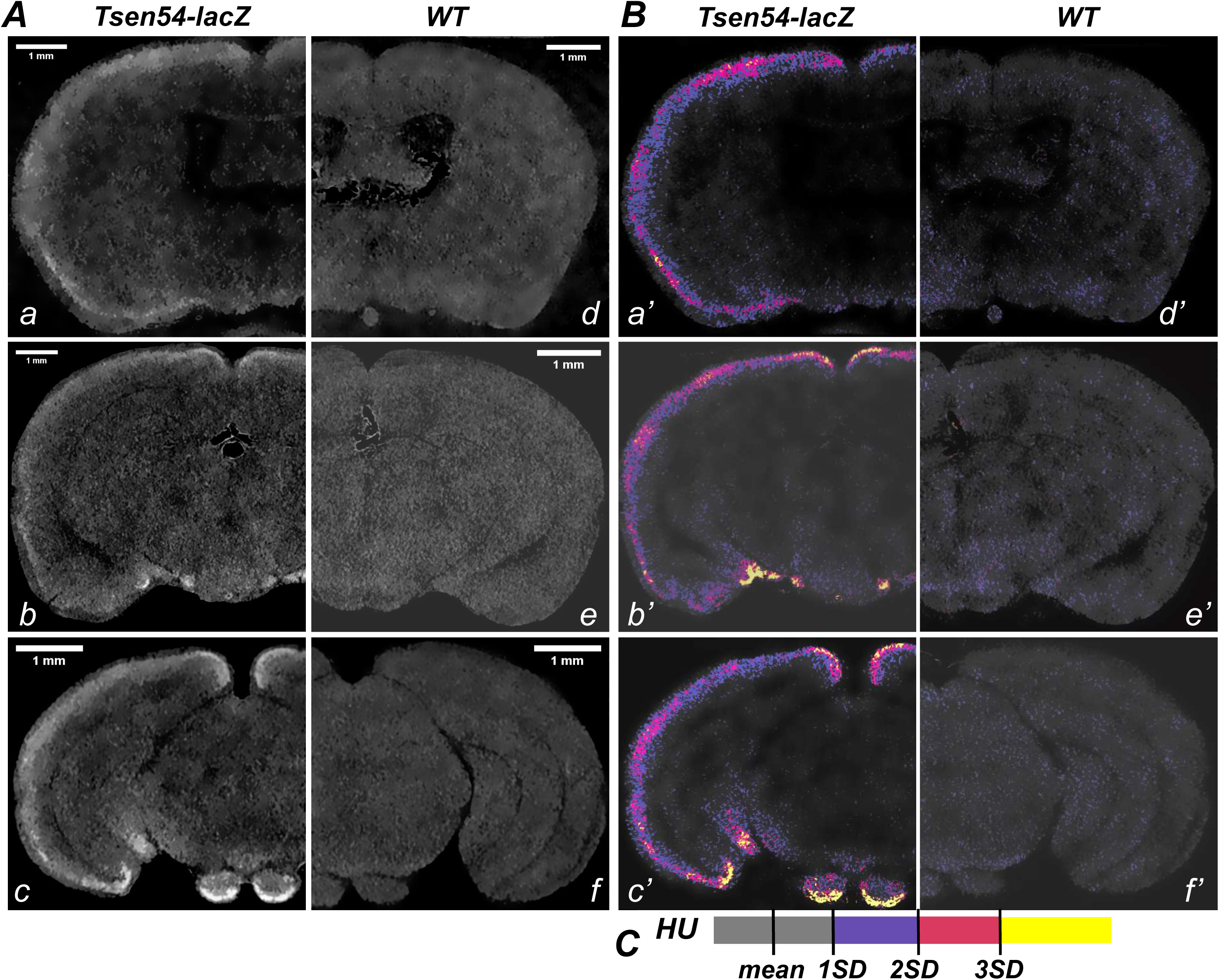
2D microCT-derived sections from the Tsen54-lacZ murine brain stained with the X-gal in presence of tetrazolium. ***A.*** 2D microCT sections of murine forebrain from *Tsen54-lacZ* (a-c) and wild type (d-f) brains. Whole-mount brain images were acquired with the resolution of 7.9 μm/voxel. ***B.*** Segmentation analysis of the microCT imaged brains were performed using CTAN (Bruker), 2D sections from *Tsen54-lacZ* (a’-c’) and wild type brains (d’-f’). HU of mean +1standart deviation (1SD) were defined as a background, while values outside of this range were colour coded: intervals between 1SD-2SD-blue; 2SD-3SD-magenta; from 3SD and up to maximum-yellow. The same values were applied for segmentation of the microCT-derived sections from wild type brains.

## Discussion

In this manuscript, we presented novel approach to analysis of *lacZ*-reporter gene expression by X-ray imaging of the whole-mount murine brain *ex vivo*. The substrate of the *lacZ/X-gal* reaction deposited *in situ* in the murine brain creates high-contrast due to the high atomic number of bromine, present in final non-soluble product of the *β-galactosidase*/X-gal biochemical reaction.

The general idea to use enzymatic activity of the reporter gene for the non-optical imaging of gene expression was previously investigated and already found broad application in medicine. Most of these studies, however, were performed using Magnetic Resonance Imaging (MRI) (50). Recent technologies allows for the detection of a reporter enzymatic activity using different probes or sensors and MRI imaging. For example, the enzymatic activity of β-*galactosidase* reporter gene can be detected in live *Xenopus laevis* embryos. β-*galactosidase* cleavage of contrasting agent (EgadMe) results in release of paramagnetic ion Gd^3+^, which, via direct interaction with water proton, increases the MR signal. While this methodology is very promising, it would be interesting to test whether EgadMe could be also used as a contrasting agent for microCT and whether this approach could be expanded to study gene expression in the *lacZ* reporter mice.

The data accumulated by IMPC consortium has demonstrated that *X-gal/lacZ* is an excellent reporter system to study gene expression in *ex vivo* murine organs. First, *X-gal* as a substrate disperses freely inside of the tissues, ensuring whole-mount staining of several organs, such as the mouse brain. Second, the intensity of the reporter *lacZ* expression faithfully reflects the activity of the promoter that drives the *lacZ* expression and is independent of the location of anatomical substructure within the brain (42). In addition, the background produced by endogenous *lacZ* activity in mouse brain previously detected biochemically (51), did not preclude from the reliable detection of reporter gene expression neither by light microscopy and does not interfere with X-ray based imaging.

An essential step that has been introduced in the protocols presented here is the sample dehydration by ethanol and paraffin embedding of the mouse brain after whole-mount *X-gal* staining. Dehydration process renders mouse brain visible to X-ray imaging without the need of any additional treatment with contrasting agents, which is instead generally mandatory in other microCT protocols developed for imaging of soft tissues. The main role played by paraffin embedding is to maintain the shape and correct morphology of both the whole sample and internal brain structures. The microCT images of the brains prepared in this way allow both to get a clear view of anatomical brain substructures and to correlate microCT-derived images with the stereotaxic coordinate of the mouse brain atlas (52). However, a substantial endogenous X-ray density due to the granular cells in the wild type brains is present. Such, the granular cell layer of cerebellum can be visualized by X-ray imaging of the dehydrated and paraffin embedded brain without the need of contrasting agents. While this observation can be instrumental for studies of the cerebellum by X-ray imaging, these background densities hamper a reliable characterization of *lacZ*-reporter gene expression in cerebellum. We circumvented these drawbacks by performing separate segmentation analysis of the cerebellum and forebrain, medulla and pons. The segmentation analysis proposed in this manuscript can be further improved by developing new advanced bioinformatics approaches to define in automatic manner the anatomic regions in which reporter gene is express.

It is important to note that imaging analysis by microCT of a brain sample does not prevent that the same sample could be afterwards thin sectioned and further investigated by light microscopy. In this way, the application of microCT imaging inverts the sequence of analytical steps classically used to obtain 3D images. In the classical approach the sample is first dissected, then imaged and at last digitally assembled as a volume image. However, this procedure often leads to a distortion of the volume information due to sample handling and make necessary the use of extensive mathematical corrections in order to avoid loss of information 53, 54). By using whole-mount microCT volume analysis instead, the whole-mount brain is first imaged by microCT, which allows for collection of maximum amount of structural information from the intact system.

Another advantage of using microCT coupled to *lacZ* reporter gene analysis is that it allows for relative quantitative comparison of gene expression level within an intact organ. This is because microCT imaging detects high-contrast products of *lacZ* enzymatic activity expressed as brightness (gray) values or Hounsfield units allowing for a relative quantitative estimation of the reporter gene expression in different brain regions. In this regard, we are aware that there is room for further improvements of our method. A greater number of analysed samples are necessary to reach an adequate level of standardization and to evaluate with good approximation the interval of linear response in the acquisition of the images by the detector used. Further, the development of digital software with improved capabilities for 3D brightness quantification will allow for a more accurate assessment of *lacZ* reaction products. If these technological requirements will be satisfied, the absolute quantification of gene expression level in reporter mouse models in the intact organs will become feasible.

Quantitative analysis of the gene expression by microCT *ex vivo* has been previously performed successfully in developing chicken embryos. The published method relies on metal-based detection with the secondary immunoreactive antibodies in developing limbs of chicken embryos (43). While the metal detection method is attractive for analysis of relatively small animals or organs, it remains to be determined whether metal antibodies detection by microCT could be applied for quantitative studies of gene expression in whole-mount organs of higher organisms like mouse. In fact, higher complexity and bigger size of the organs of mouse could obstruct the accessibility of recognition sites to antibodies in whole-mount organ preparations. In addition, metal immunodetection method suffers from possible non-specific deposition of antibodies especially in folded complex biological samples and suffered from unspecific epitope recognition by antibodies (43). Our method, instead, does not require specific probes or antibodies for every specific gene tested because it uses *lacZ* as a common gene reporter. Furthermore, another advantage of *lacZ* as a substrate for gene expression studies by microCT is that it relies on commonly available and low-cost reagents.

The microCT gene expression analysis based on the *lacZ* reporter can be easily integrated into existent high throughput phenotyping efforts for gene expression analysis currently performed in highly specialized and technologically advanced centres such as Allen Brain Institute and IMPC consortium. A bank of the *X-gal/lacZ* stained tissues and organs was already generated by IMPC consortium (www.mousephenotype.org) (42) and all those murine lines represents available candidates for microCT based studies. In addition, because our method does not require histological tissue sectioning and challenging 3D reconstructions, it can be performed in any laboratory equipped with a laboratory microCT-imaging device.

Here we conducted experiments using top-bench Bruker Skyscan 1272 microCT, which allows the analysis of the murine brain with resolution ranging, in our experiments, from 7 to 20 microns. This resolution while allows to detect the anatomical substructure in which the reporter gene is expressed, it does not allow to characterize the nature of the cells expressing the gene. This limitation could be overcome by using advanced microCT machines such as nanoCT systems, propagation-based X-ray phase-contrast tomography or synchrotron based X-ray microtomography allowing substantially increase the imaging resolution. Several groups demonstrated the feasibility of these technologies for analysis of murine organs and specifically mouse brain at cellular resolution (55, 56; 57, 58,59). Application of these advanced X-ray based technologies will allow to improve the resolution of our X-ray based gene expression analysis method to the cellular level.

We have characterized patterns and relative expression levels of the Tsen54 gene in different murine brain regions. *Tsen54-lacZ* reporter is diffusely expressed in the cortex with the highest expression observed in the Retrosplenial Cortex (RSC). Both in humans and in rodents the RSC is important for a variety of cognitive tasks including memory, navigation, and prospective thinking (60, 61). In rodents, it is also critical for spatial memory such as allocentric working memory, and functions in which animals have to detect if a spatial and arrangement is novel or familiar (62). The high expression of *Tsen54* gene in the RSC suggests the role *Tsen54* may play for the cognitive processes and specifically in formation of spatial memory in human and mice.

Our microCT based analysis also demonstrated high*Tsen54* expression in the temporal lobe of dentate gyrus. Histological analysis shows that the *Tsen54* expression is limited to the granular cell layer of dentate gyrus. The dentate gyrus is a substructure of the temporal lobe region, which is composed of anatomically related structures (CA fields, dentate gyrus, and subicular complex) that are essential for declarative memory (conscious memory for facts and events) and the spatial memories in humans and in rodents (63). This result might suggests that *Tsen54* gene as a part of TSEN complex play an important role in the regular functions of this brain region. *Tsen54* gene controlling protein synthesis via tRNA splicing process might be important for the memory formation.

Lateral mammillary nucleus is a part of hypotalamic system of interbrain is also characterized by high *Tsen54-lacZ* expression. Mammillary nuclei is an anatomical substructure of the mammillary bodies composed of the medial and lateral nuclei. This two subregions are characterized by very different functions. Such that the medial mammillary bodies via inputs from the ventral tegmental nucleus of Gudden support memory processes. In parallel, the lateral mammillary bodies, via their connections with the dorsal tegmental nucleus of Gudden, are critical for generating head-direction signals (rev in 64). This result indicates specific role of Tsen54 in processing of the signals involved in head-direction movements.

Here we have also observed that *Tsen54-lacZ* have high expression in cortical plate of the murine brain. The subregions with the highe expression forms fork-like structure at rostral part of the brain, which is composed of several substructures: anterior olfactory nucleus, bed nucleus of the accessory olfactory (BA), the nucleus of the lateral olfactory tract (NLOT), with the lateral preoptic area tract (LPO) and by the vascular organ of the lamina terminals. It would be interesting to understand why these regions has specific requirements for the high *Tsen54* expression. Further studies using murine model with the conditional deletion of *Tsen54* gene will allow to address this question.

Especially interesting is an expression pattern observed in hindbrain. The particularly strong signals have been observed in the pons, with the highest expression limited to pontine gray. This is an interesting observation, because in PCH patients with the *TSEN54* mutations noticeable pons flattening of several degrees is observed (36), suggesting an autonomous role of the *Tsen54* gene in pones development and functions. In medulla, high expression in precerebellar nuclei: external cuneate, lateral reticular and spinal trigeminal is observed. This pons and medulla substructures are important for proper signal exchange with cerebellum exerting the fine control on balance, motor movements and motor learning (65). High *Tsen54* expression marks the nuclei, listed above, from which excitatory mossy fibers originate, suggesting possible role of Tsen54 gene in normal transmission of excitatory signal. The mossy fibers synapse on cerebellar granule cells providing excitatory input to Purkinje cells, which in turn are inhibitory neurons that control the main cerebellar output system (66). The loss of equilibrium between the excitatory and inhibitory signals in cerebellum might be a reason for the neuronal degeneration and death as observed in cerebellar hypoplasia affected individuals. Indeed, histological analysis demonstrated *Tsen54* in the cerebellar stellate cells of molecular layer and in the Purkinje cells as well. It is interesting to note that in PCH patients carrying mutation in *TSEN54* gene show almost the complete loss of the Purkinje cells, suggesting that these cells might be specifically targeted by *TSEN54* gene dysfunction leading to their cell death. Therefore, it would be important to understand if TSEN54 plays an autonomous role in Purkinje cell degeneration or it happen due to the insufficient signal, which arrived to the Purkinje cells from other regions, which are highly expressing Tsen54 gene. All these possibilities could be addressed in further studies using murine models of PCH with the loss of *Tsen54* gene function.

In summary, here we identified a number of specific brain regions and substructures with high *Tsen54* gene expression, which might suggest a strong functional dependence on *Tsen54* gene function. Our follow up studies will be focused on the mechanisms by which *Tsen54* gene executes its role in neurogenesis and explain why mutations in this gene lead to the neurodegeneration in the form of PHC.

Overall here we have presented the novel method of three-dimensional characterization of lacZ reporter gene expression by X-ray microCT imaging and illustrated its applicability to analysis of the gene expression pattern in intact murine brain. The simplicity of the *lacZ/X-gal* staining protocol and availability of the reporter animals for virtually any gene of mouse genome would allow for the high-throughput application of described methodology.

## Material and Methods

### Ethics Statement

After weaning, mice were housed by litter of the same sex, 3 to 5 per cage and maintained in a temperature-controlled room at 21 ± 2 °C, on a 12-h light-dark cycle (lights on at 7AM and off at 7PM), with food and water available *ad libitum* in a specific pathogen-free facility. The experimental protocols and animal care procedures was reviewed and approved by the Ethical and Scientific Commission of Veterinary Health and Welfare Department of the Italian Ministry of Health (protocol approval reference: 118/2012-B), according to the ethical and safety rules and guidelines for the use of animals in biomedical research provided by the Italian laws and regulations, in application of the relevant European Union’s directives (n. 86/609/EEC and 2010/63/EU).

### Animals

*Tsen54^tm1b^/+* heterozygous animals were derived from ES cells obtained from IKMC consortium. *Tsen54^tm1a(EUCOMM)Wtsi^* (https://www.mousephenotype.org/data/genes/MGI:1923515), The *Tsen54-lacZ* reporter allele was engineered by the genetic modification protocol developed by IKMC consortium and the targeting strategy is deposited on the IMPC portal https://www.mousephenotype.org. In brief knockout mice were generated from *Tsen54tm1a*(EUCOMM)Wtsi ES cells produced (Figure 1A). The animals carrying *Tsen54tm1a*(EUCOMM)Wtsi were breed with the with *ROSA26Cre* (MGI: 5285392) animals as a result heterozygous animals with the Tsen54Tm1b allele were produced. *Tsen54Tm1b* allele is a knockout of *Tsen54* gene in which exon 6 is replaced by *lacZ* reporter while neo cassette is removed from the targeted locus (Figure 1A). We have previously demonstrated that *Tsen54* gene is essential for the embryonic development and homozygous *Tsen54* knockout embryos do not developed past implantation (29). We have observed that heterozygous *Tsen54^Tm1b/+^* animals have a normal lifespan. We did not observed any gross anatomical, physiological abnormalities in *Tsen54^Tm1b/+^*. animals (data not shown)

### Whole mount mouse brain preparation for a microCT imaging

All solutions used for the brains preparations were filtered with 0.4 μm filtering system. Adult mice (4-12 weeks old) were anaesthetized with 2.5% avertin solution (100μl/10g of body weight) and transcardially perfused, first with PBS (5 ml per animal) and then with 4% PFA (20 ml per animal). Brain were extracted from the scalp and fixed for additional 30 min with 4% PFA on ice and then were incubated for 30 min in wash buffer (2mM MgCl_2_, 0.01% sodium deoxycholate, 0.02% of Nonidet-P40, 0.1M sodium phosphate buffer pH 7.3) on ice. Brains were stained in *X-gal* staining solution containing 1mg/ml of *X-gal* stock solution (25mg/ml*X-gal* (Sigma-Aldrich, B4252) in dimethylformamide), 0.2% of potassium ferrocyanide (Sigma-Aldrich, P-9387) and 0.16% potassium ferricyanide (Sigma-Aldrich,P-8131) in wash buffer for about 48 hours at 37°C. Staining with the tetranitroblue tetrazolium chloride was performed as described above, except that potassium ferrocyanide and potassium ferricyanide was replaced with the tetranitroblue tetrazolium chloride (Sigma cat:87961) at final concentration 3,3 μg/ml. After staining completed blue precipitates might be visually apparent. Brains were washed in wash buffer and additionally fixed with 4% PFA overnight in the cold room and embedded in paraffin as described (67). Dehydration was performed by sequential changes of ethanol: 50% ethanol for 2 hours, 70% ethanol overnight, 95% ethanol for 30 min., all treatments were performed in the cold room, brain then were incubated in 100% ethanol at room temperature for 2 hours and transferred into xylene for 45 min. Xylene then was replaced with the xylene/paraffin solution mixed at 1:1 ration and brains were incubated at 56^0^C for 45 min., then paraffin/xylene solution was replaced by the paraffin and samples were incubated overnight at 56^0^C. Next morning paraffin was replaced with the fresh paraffin and brains were incubated for 1 hour at 56^0^C, transferred in the tissue embedding molds and allowed to polymerize for about 1 hour. Paraffin then was trimmed around the brain to feat into the holder of the microCT scanning machine.

### Immunohistochemistry

The mouse brains were *X-gal/FeCN* stained, embedded in paraffin and sectioned at 16μM using Leica RM 2135 Manual Microtome. The sections were deparaffinized in xylene, immersed in decreasing concentrations of ethanol, and rehydrated in water. All sections for immunostaining were processed for microwave-enhanced antigen retrieval in 0.01 M sodium citrate buffer pH 6.0 for 10 min in a 700-W microwave oven at maximum power for 10 min. Endogenous peroxidase activity was blocked with 1% H2O2 in water for 30 min in the dark then rinsed in PBS/0.1% Tween solution. Blocking was performed in a PBS solution containing 5% donkey serum (D9663; Sigma-Aldrich) for 60 min at room temperature. The sections were stained with Anti-beta Galactosidase antibody (ab9361; Abcam) diluted 1:500, overnight. After several rinses in PBS/0.1% Tween solution the sections were incubated with Biotinylated Goat Anti-Chicken IgY (AVESLAB) antibody for 1h in PBS-Serum, rinsed with PBS/0.1% Tween and incubated with the Vectastain Elite ABC reagent for 30 min (PK-6100 Vectastain Elite ABC Kit; Vector). The excess of ABC regent were washed with PBS and the reaction was developed by DAB (3,3′-Diaminobenzidine tetrahydrochloride (D5905-50TAB; Sigma-Aldrich) and 0.01% H2O2 solution for 5 min washed with 5mM EDTA in PBS and water. The immunostained sections were dehydrated and mounted using Eukitt (cat 09-00100, Bio Optica) mounting medium and visualized using a motorized LMD 7000 microscope (Leica Microsystems).

### Light microscopy

*X-gal*/*FeCN* stained brains, embedded in paraffin, were sectioned at 16μm and stained with eosinY 1% aqueous solution (Bio-Optica 05-M10002). Images were obtained with the stereomicroscope MZ12 (Leica) equipped with color camera.

### XPS analysis of in vitro b-galactosidase reaction products

Reaction was performed with 0.16 units of b-galactosidase (Sigma cat: G4155) or without enzyme for the control *in X-gal* staining solution containing 1mg/ml of *X-gal* stock solution (25mg/ml*X-gal* (Sigma-Aldrich, B4252) in dimethylformamide), 0.2% of potassium ferrocyanide (Sigma-Aldrich, P-9387) and 0.16% potassium o ferricyanide (Sigma-Aldrich,P-8131) in wash buffer for about 48 hours at 37°C in 50 μl. The entire volumes of the reaction and the control were deposited on the glass as a thin film and dried at room temperature. XPS measurements were carried out in a spectrometer Escalab MkII (VG Scientific Ltd, UK) by using Al Kα source. The acquisition parameters were the following: analyser pass energy = 40 eV, large area lens mode A1 x 22 with an analysis diameter of about 10 mm.

### MicroCT imaging

Brain samples were fixed using Orthodontic Tray Wax (09246 Kerr) and 3D images were performed by Skyscan 1172G (Bruker, Kontich – Belgium) using a L7901-20 Microfocus X-ray Source (Hamamatsu). The source uses an X-ray tube with beryllium output window (150 µm thick), with a focal spot size of 7 µm (10W) or 5 µm (at 4W) capable of operating at a maximum tube voltage of 100 kV. The X-ray tube voltage adjustable range is 20kV to 100kV and was set at 39 kV; the X-ray tube current adjustable range is 0 µA to 250 µA and was set at 240 µA (9W).

Scanning parameters (Skyscan 1172 MicroCT Control Software):

- No Filter
- Near Camera Position
- Dim. 1000×666 (1K)
- Large Pixel Size
- Exposure time 60 ms
- Magnification 20 µm
- Rotation step 0.4°
- Partial width 88%

Single images (stored as TIFF-file) were reconstructed from projection images taken every 0.4° of rotation over 360°, between 650 and 700 projections and total scanning times were approximately 30 min.

Projection

Reconstruction Parameters (NRecon Skyscan Reconstruction Software):

- Smoothing 2
- Misalignement compensation 25
- Ring artefact reduction 6
- Beam-hardering correction 40%
- Smoothing kernel Gaussian

Total reconstruction times were approximately 5 min.

The reconstructed tomographic dataset was stored as .BMP(8) -files (dim. 1000×1000) and read into 3D Visualization Software CTVOL v. 2.0 (Bruker) to make the volume rendering views and movies.

The density quantifications were calculated using Analyser Software CTAN v. 1.13 (Bruker) and CTVOL 2.0.

## Supplementary information

**Movie 1.**
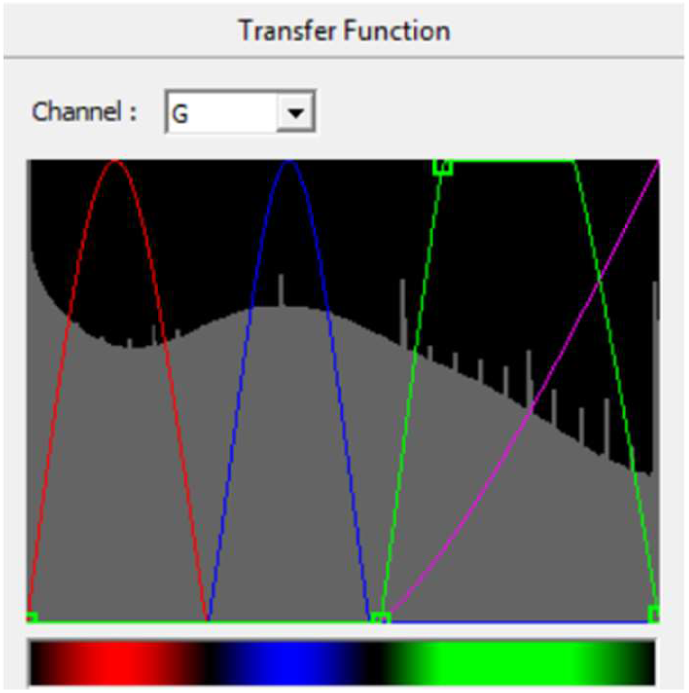
MicroCT derived RGB volume of the *Tsen54-lacZ* mouse brain stained with *X-gal/FeCN* method. The volume image was acquired using camera resolution of 1K (image matrix of 1000×575 pixel with the 7,9 μm/voxel size). Screenshot of the transfer function editor windows of the CTVOX analyser (Bruker software) demonstrates setting of the RGB transfer function curves for building a color volume-rendered 3D model; color coded for the tissue density function: red-blue-green; transparency level defined by the purple line.

**Movie 2.**
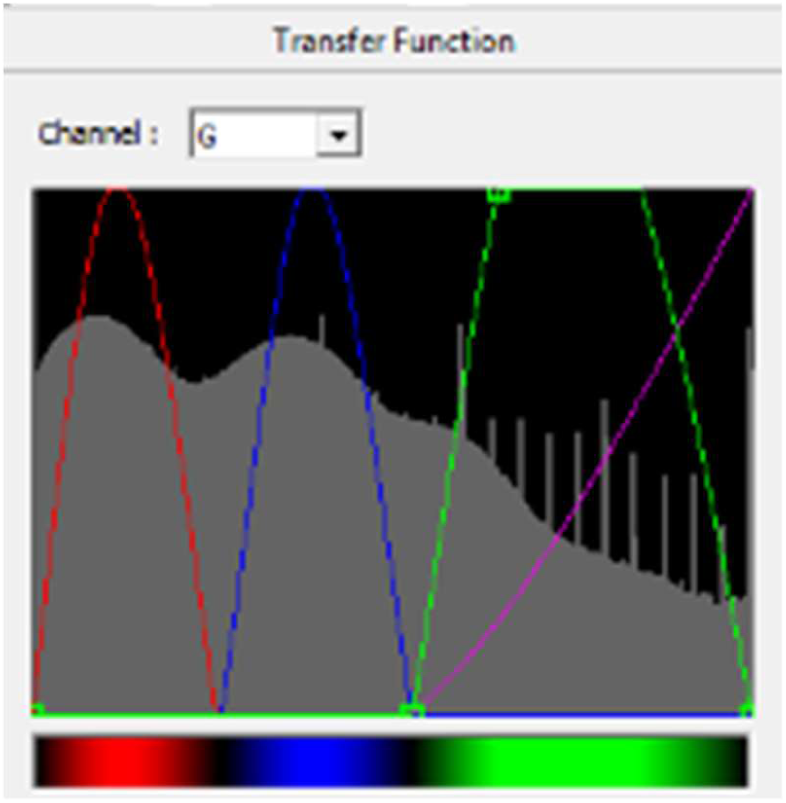
MicroCT derived RGB movie of the wild type mouse brain stained with X-gal/FeCN method. The volume image was acquired using camera resolution of 1K (image matrix of 1000575 pixel with the 7,9 μm/voxel size). Screenshot of the transfer function editor windows of the CTVOX analyser (Bruker software) demonstrates setting of the RGB transfer function curves for building a color volume-rendered 3D model; color coded for the tissue density function: red-blue-green; transparency level defined by the purple line.

***Movie 3:*** Segmantation analysis of the *Tsen54-lacZ* brain, whole-mount stained with *X-gal/FeCN* and imaged by microCT. MicroCT imaging was performed with the resolution of 20 μm of voxel size. Calculated using CTAN (Bruker software) mean of Hounsfield units from the *Tsen54-lacZ* brain and one standard deviation (1SD) were defined as a background, while values outside of this range were colour coded: values from 1SD to 2SD-blue; from 2SD-3SD-magenta; from 3SD and up to maximum – yellow. The reconstruction of the segmented *Tsen54-lacZ* brain was performed using CTVOL analysis (Bruker software).

***Movie 4***: Virtually reconstructed in three-dimension expression pattern of *Tsen54* gene. The reconstruction was based on the microCT analysis of the *Tsen54-lacZ* mouse brain *ex vivo* and performed using A. Brain Atlas BrainExplorer2 software using Allen Mouse Common Coordinate Framework option. The selected for the 3D reconstruction brain structures are listed in Table 1. Dentate gyrus was not included because expression was observed only in the temporal lobe of dentate gyrus. Cerebellum also was excluded from virtual reconstruction for the best visualization of the nuclei of the nuclei within medulla.

***Movie 5*.** Segmantation analysis of the *Tsen54-lacZ* brain whole-mount stained with *X-gal/Tetrazolium* protocol and imaged by microCT with the resolution of 20μm of voxel size. Calculated using CTAN (Bruker software) mean of Hounsfield units of the *Tsen54-lacZ* brain and one standard deviation (1SD) were defined as a background, while values outside of this range were colour coded: values from 1SD to 2SD-blue; from 2SD-3SD-magenta; from 3SD and up to maximum – yellow. The reconstruction of the segmented *Tsen54-lacZ* brain was performed using CTVOL analysis (Bruker software).

## Acknowledgements

We are grateful to S. Manghisi, A. Grop, E. Manghisi, G. D’Erasmo, M. Parmigiani and S. Gozzi from European Mouse Model Archives (EMMA), Monterotondo, Italy for excellent technical support in animal handling, sample preparation and data analysis. We are grateful to Dr. G. D. Tocchini-Valentini (IBCN, CNR, Monterotondo, Italy) for guidance in X-ray based methods of imaging and to Dr. D. Marazziti (IBCN, CNR, Monterotondo, Italy) for critical reading of the manuscript. Special acknowledgments to A. Ferrara and T. Cuccurullo for secretarial work.

